# Internal and external features of wild birds after collisions without apparent trauma in Japan

**DOI:** 10.1101/2021.05.25.445584

**Authors:** Nana Ushine, Aki Tanaka, Tatsuo Sato, Masaki Nonagase, Shin-ichi Hayama

## Abstract

Wild birds often require rehabilitation after collisions, even with no apparent injury. Information about aftermath of collisions is still scares. Here, we investigated external characteristics and clinical features of the internal organs of wild birds that experienced collision and compared them with birds admitted to rehabilitation center for other reasons. Necropsy was performed on 55 bird carcasses from Passeriformes and Coraciiform. Five external characteristics were recorded before necropsy including; cause of admission, keel score, life stage, fat score, and number of days before death. The median survival time was calculated by Kaplan-Meier estimates. Data on external and internal features were compared using univariate and multivariate multinomial regressions. There was no significant difference in the median survival time among the causes of admission: 1 day for collision, 2 days for trauma, and 2 days for malnutrition. Kidney discoloration was more significantly associated with collision than with other trauma (p = 0.01). Although no apparent kidney abnormality (including enlargement) were observed, anterior lobe was significantly larger than posterior lobe with collision compared with malnutrition (p = 0.045). Birds that experienced collision exhibited a higher fat score than malnourished birds (p = 0.03). Our results suggested that wild birds with abundant fat were more likely to be admitted due to collision. The gross characteristics of collision included kidney discoloration and anterior lobe extension, which was possibly due to rupture of renal blood vessels by blunt external force. From these findings, it was considered that collision caused major axis of anterior lobe significantly larger than posterior lobe, even though no abnormal finding in renal size such as hypertrophy was recognized. This was the first study to evaluate the cause of admission, necropsy results, and external characteristics in wild birds admitted to rehabilitation centers. Absolute cage rest should be adhered to restore renal function for those birds admitted due to collision, and handling and treatment should be minimized to avoid excess movement of the birds.

## Introduction

Wild animals experience a variety of anthropogenic traumas [1]. In the United States, there were approximately one billion anthropogenic-related wild bird deaths annually, and approximately 60% of rehabilitation cases result from collisions with buildings that caused no visible trauma [2]. In France, 76% of the birds in rehabilitation were admitted for collision accidents without apparent trauma, including falls from nests [3]. Similar numbers had been reported in Japan [4-6], with collision without trauma representing over 70% of rehabilitation cases [7, 8]. Wild birds accounted for 90% of all wildlife rehabilitation cases in Japan [9, 10]. The number of cases attributed to collision had increased globally with the development of urban structures [11]. Birds that experience trauma due to collision and were brought to rehabilitation centers generally had low recovery rates and are less likely to be released back into the wild [12, 13]. Previous research was focused primarily on designing structures to prevent collision accidents, and only a few studies have investigated the association between the cause of admission to rehabilitation centers and the injuries sustained [14-17]. In human medicine, the degree of damage to internal organs due to collision was related to time of clinical assessment, and the early management of damaged organs improves prognosis [18]. In veterinary medicine, the low recovery rates of birds rehabilitated after collision suggested that this type of accident significantly affect the vital organs [19, 20]. Therefore, in this study, we aimed to investigate the external characteristics and clinical features of the internal organs of wild birds that experienced collision, and compared them with other admission causes.

## Materials and methods

Fifty-five wild passeriform and coraciiform birds were examined that died at the Wild Bird Rehabilitation Center, Chiba Prefecture, Japan, from June 2016 to August 2020. Necropsies were performed at the Nippon Veterinary and Life Science University, Tokyo, Japan.

The following data were recorded at the time of admission: causes of admission, five-grade keel score, fat score [21], life stage, and number of days to death. Life stage was determined by plumage in both orders, that is, birds with complete adult plumage were defined as “juveniles and older,” and birds without such a plumage were defined as being “younger than juveniles.” Additionally, life stage was determined by skull pneumatization in the passeriform specimens. The following body parts were evaluated during necropsy: beak, nostril, abdomen, wings, intraoral cavity, neck, back, waist, and legs. Moreover, lungs, trachea, heart, liver, skull, body cavity, stool, kidneys, and anterior to posterior kidney lobe ratio were examined. Abnormal findings at the time of necropsies were recorded. Each organ was cut along the sagittal plane, and macroscopic examinations were performed.

Kaplan–Meier analysis and log-rank tests were used to compute a 50% survival analysis, with the median survival period associated with the cause of admission. Life table analysis was performed to analyze the median survival time with a survival probability of 50%, and a 95% confidence interval (CI) when 50% survival probability was not attained by the end of the study. Univariate and multivariate multinomial logistic regression analyses were used to examine associations among causes of admission, life stage, and necropsy findings. The results were reported as odds ratios and 95% CIs. The life stage and necropsy indices were defined as independent variables, and the three causes of admission were defined as dependent variables. The variables with p < 0.1 in the univariate multinomial logistic regression and those considered clinically important were included in the multivariate analysis to construct the final model. For the statistical estimation and inferences, two-sided hypothesis tests were performed with a 5% significance level. Stata/IC 14 statistical software (Stata Corp. LP, College Station, TX, USA) was used for all analyses.

## RESULTS

The species and number of rescued birds were as follows: *Passer montanus* (*n* = 40), *Parus minor* (*n* = 4), *Hypsipetes amaurotis* (*n* = 2), and *Sturnus cineraceus* (*n* = 6) belonging to Passeriformes; and *Alcedo atthis* (*n* = 3) belonging to Coraciiformes. The external characteristics of the 55 birds and necropsy findings were shown in Table 1. Kidney discoloration was recognized for the first time in the present study.

**Table 1.**
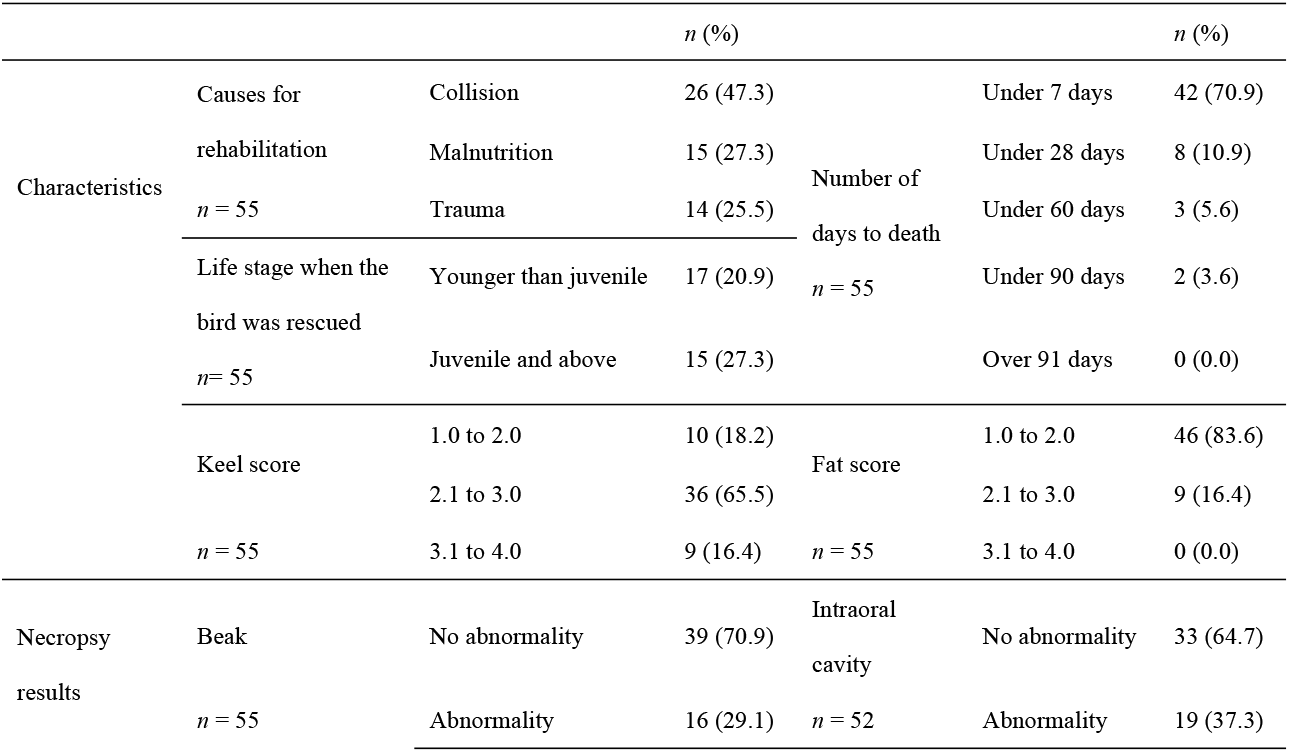
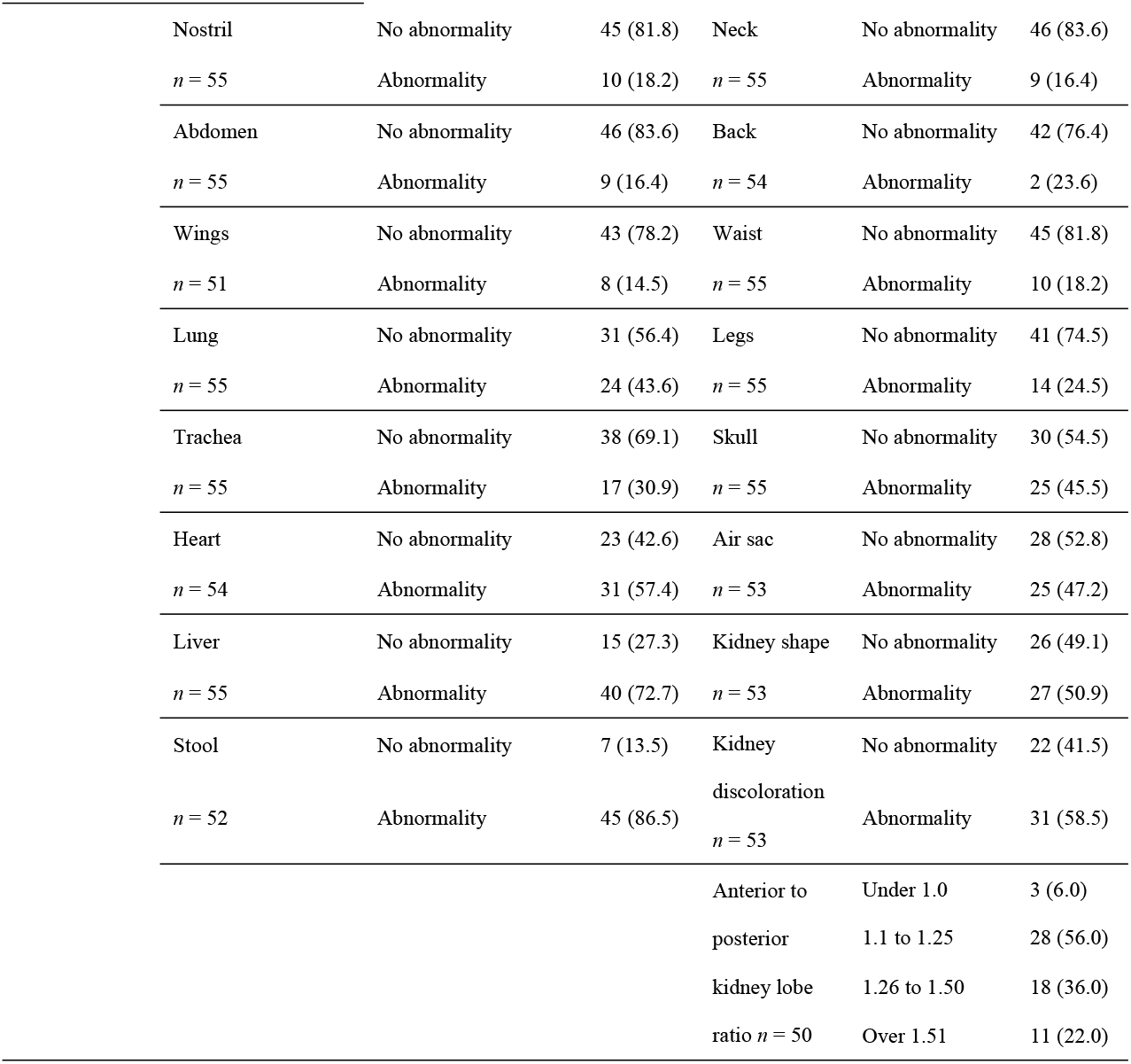
Characteristics of the 55 birds investigated in this study from June 2016 to August 2020 in Chiba Prefecture, Japan.

There was no significant difference in the survival time among the three causes of admission (collision, trauma, and malnutrition; Fig. 1). The median survival time was 1 day for collision at 50% survival probability with a 95% CI of 30–67, 2 days for trauma at 50% survival probability with a 95% CI of 24–71, and 2 days for malnutrition at 50% survival probability with a 95% CI of 23–72.

**Figure 1.**
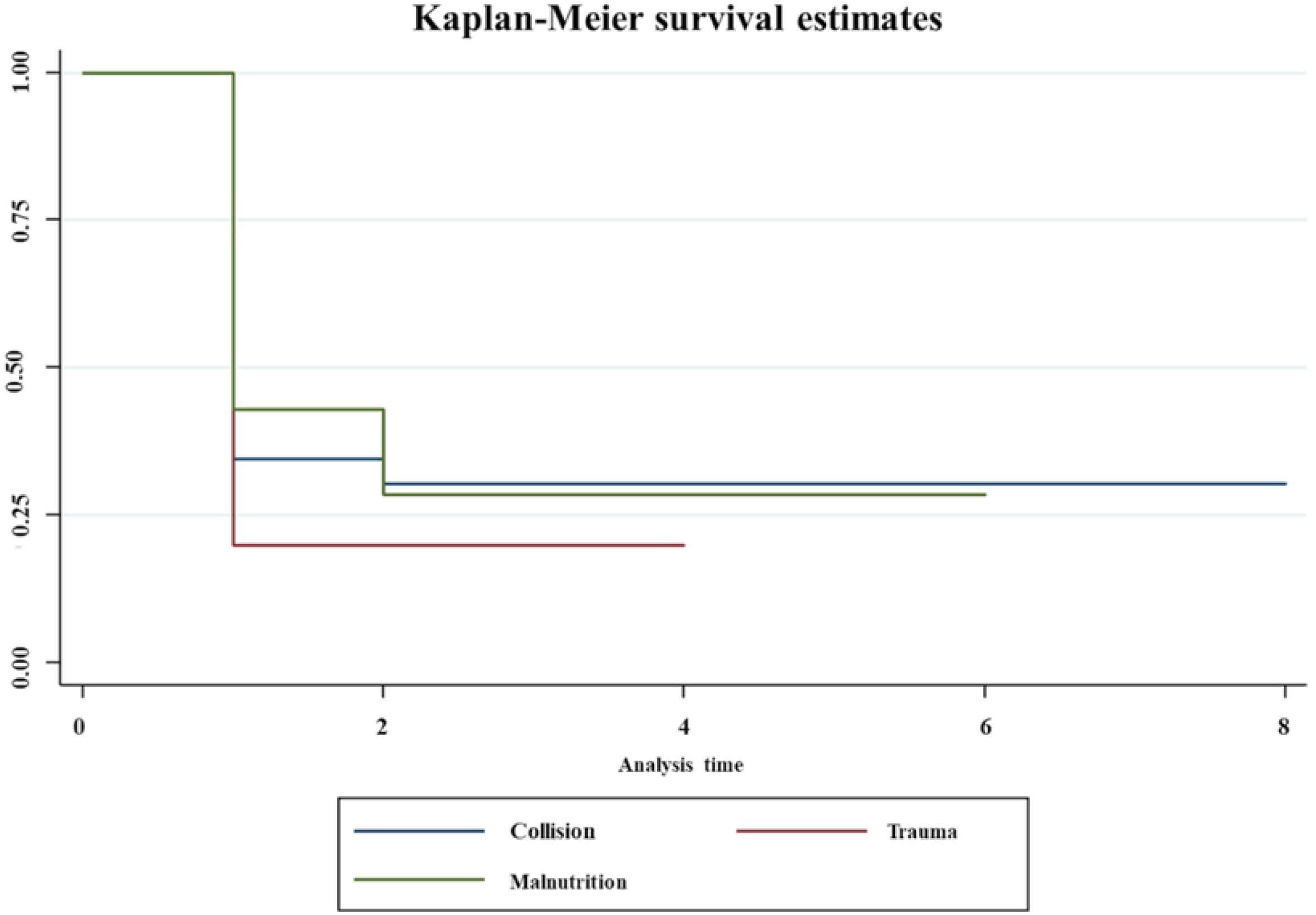
Kaplan-Meier plot showing time to death caused by collision, trauma, and malnutrition in 55 rescued wild birds in Chiba Prefecture, Japan from June 2016 to August 2020. There was no significant difference in survival time between causes for rehabilitation (p = 0.52).

The results of the univariate analysis were shown in Table 2. Compared with collision, trauma was associated with fat score (95% CI 0.12–1.12; p = 0.08), lung abnormalities (95% CI 1.50–25.47; p = 0.01), kidney discoloration (95% CI 0.02–0.36; p = 0.001), and anterior to posterior kidney lobe ratio (95% CI 0.00–0.54; p = 0.02). Additionally, malnutrition was associated with life stage (95% CI 0.19–0.94; p = 0.02), fat score (95% CI 0.03–0.44; p = 0.002), keel score (95% CI 0.09–1.13; p = 0.08), back abnormalities (95% CI 1.00–17.57; p = 0.05), leg abnormalities (95% CI 0.92–18.52; p = 0.06), lung abnormalities (95% CI 1.02–16.00; p = 0.046), and anterior to posterior kidney lobe ratio (95% CI 0.00–1.03; p = 0.045) compared with collision.

**Table 2.**
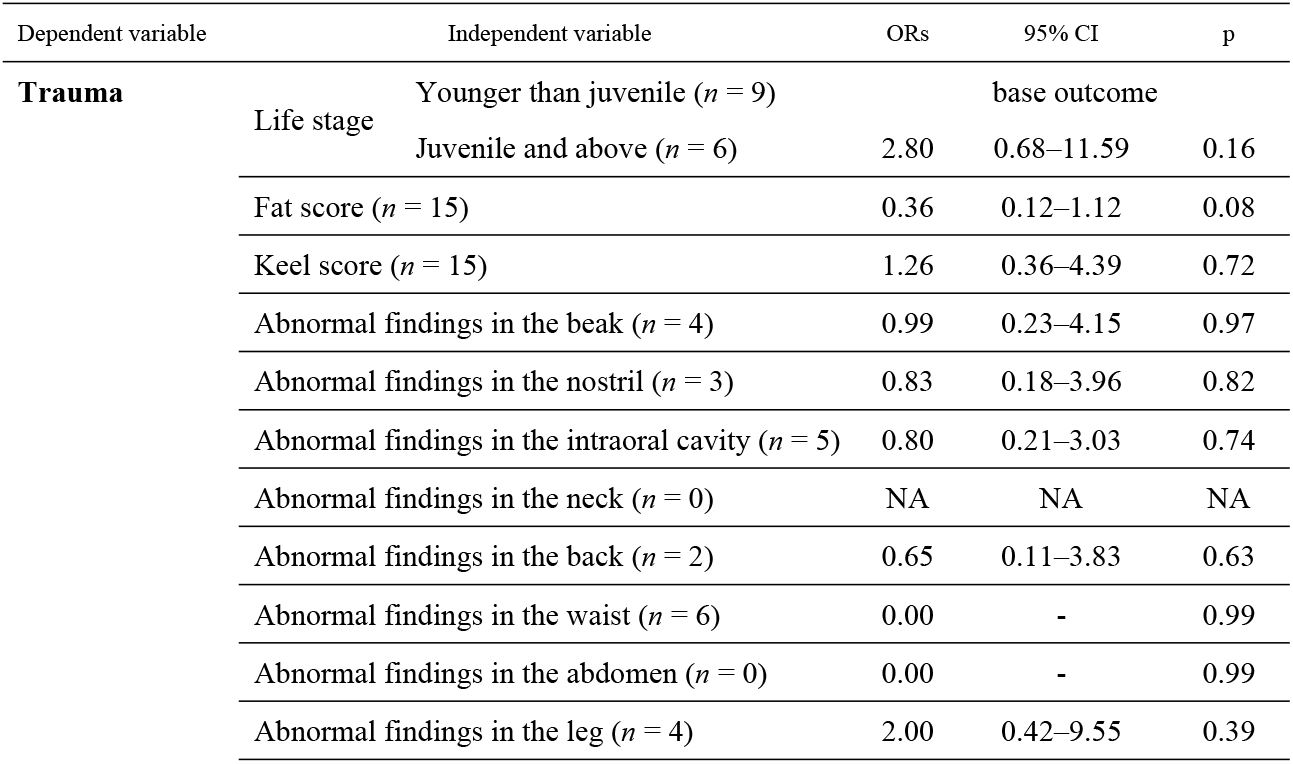
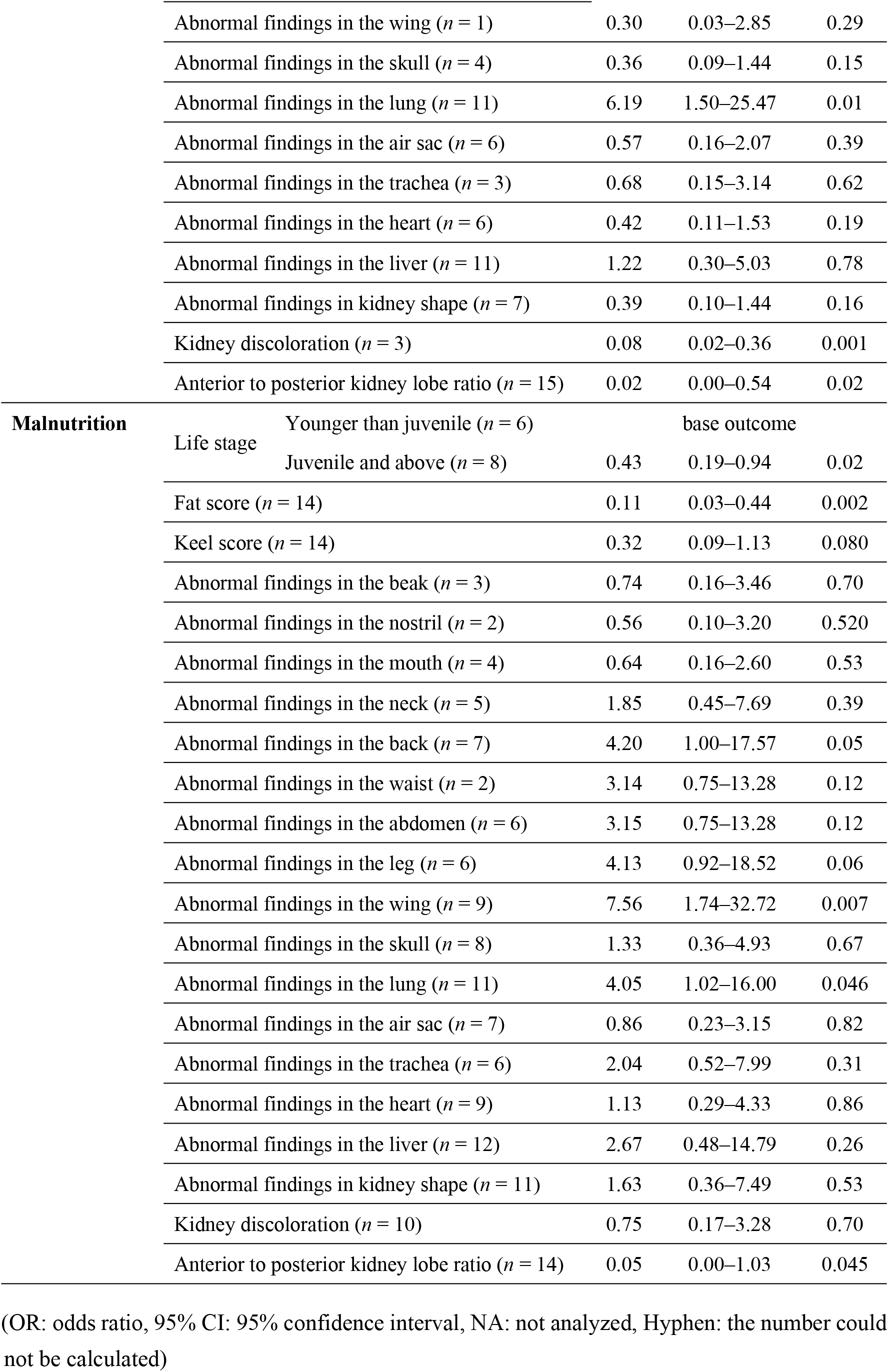
Univariate multinomial logistic regression analysis of the association of the cause of admission and life stage fat score, keel score, and necropsy findings in 55 wild birds admitted in Chiba Prefecture, Japan from June 2016 to August 2020.

The multivariate analysis showed that collision was less likely to cause abnormalities in the lung and more likely to cause a kidney discoloration. Additionally, collision resulted in a higher anterior to posterior lobe ratio than trauma (p = 0.04; Table 3), and was associated with a higher fat score and anterior to posterior kidney lobe ratio than malnutrition. Lung abnormalities occurred less frequently from collision or trauma than from malnutrition (p = 0.02; Table 3).

**Table 3.**
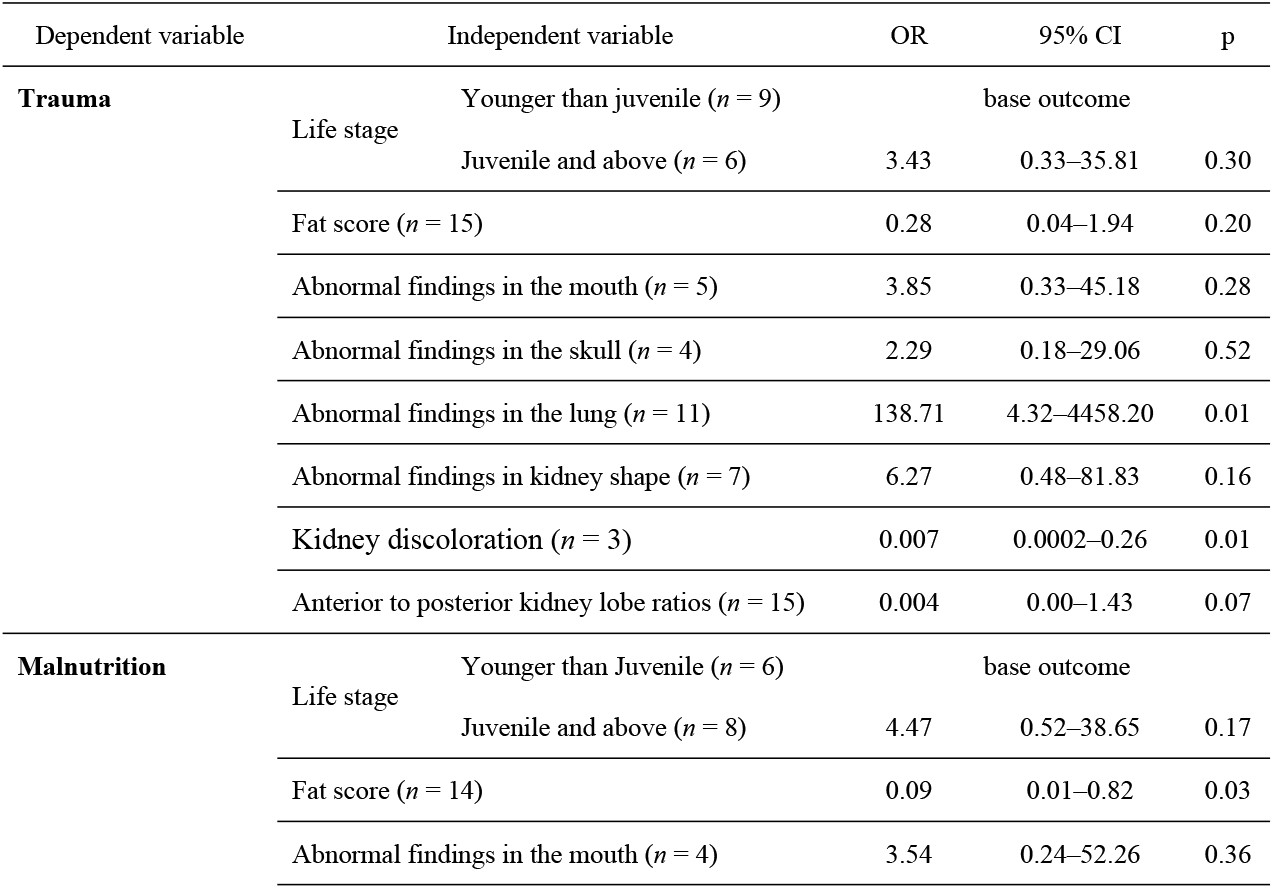
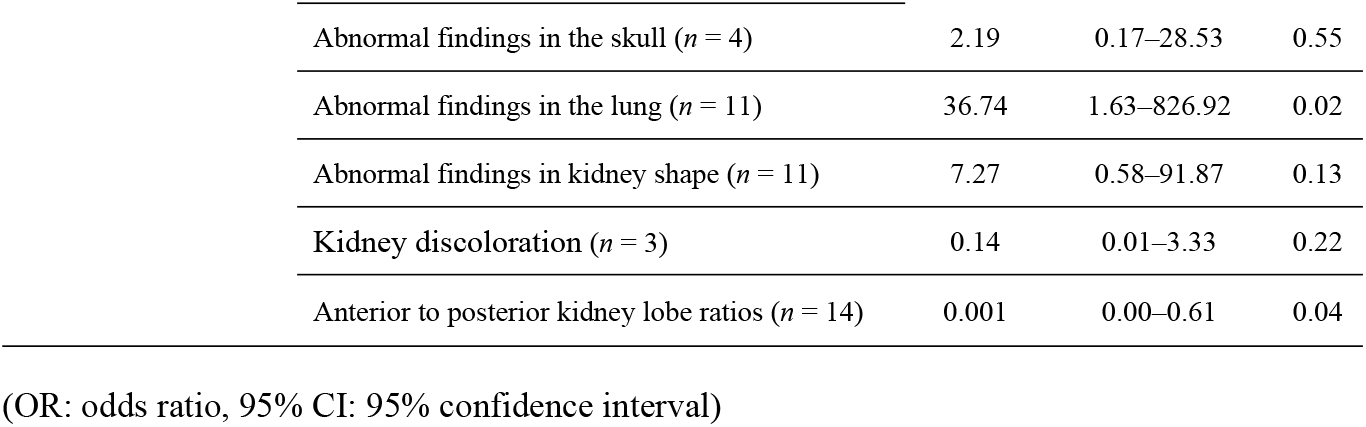
The final model for the multivariate multinomial logistic regression analysis of causes of admission in 55 wild birds admitted in Chiba Prefecture, Japan from June 2016 to August 2020.

## DISCUSSION

Our study indicated a significant association between collision and anomalies in the anterior to posterior kidney lobe ratio in accidents involving physical damage. A bird admitted to the Wild Bird Rehabilitation Center because of a collision was more likely to have a higher score for fat than one admitted for malnutrition. The necropsy procedure was based on previous reports [22-24]. The present report was the first to compare necropsy findings of collision accidents and other types of causes of admission in wild birds. Life stage was considered a confounding factor in this analysis because a collision in nestlings was typically the result of falling from the nest. Unlike a collision with a building during flight, nestling collisions may affect the posterior body. Due to the small sample size, a significant association was not recognized between life stages, the external characteristics, and internal organs features of wild birds admitted because of collisions compared with other causes.

To the best of our knowledge, there were no previous reports in the literature on renal hypertrophy that demonstrated the relative sizes of the anterior and posterior kidney lobes, associating them with the cause of death. In avian anatomy, the anterior lobe is innervated by the nerve plexus that regulates the thigh; damage or extension to this anterior lobe can cause posterior lobe insufficiency and severely affect the prognosis of rescued birds [25]. Rhabdomyolysis, a cause of capture myopathy in wildlife [26], caused a large amount of myoglobin to flow into the kidney, which damages the glomerulus and renal tubules and results in kidney swelling and discoloration as macroscopic findings [27]. A collision may damage the skeletal muscle similarly to capture myopathy. Therefore, it is possible that both situations cause a micropathological condition in the kidney. In this study using a micropathological examination, only the anterior kidney lobe was confirmed to be extension. From these findings, it was considered that collision impact might have extended major axis of anterior lobe compared to posterior lobe, even though there was no notable change in the size of kidney. Anterior kidney lobes are exposed and located outside the pelvis, unlike the middle and posterior kidney lobes which remain inside the pelvic structure, therefore, anterior kidney lobes are subjected to impact from collision.

This study reported the impact of minor collisions on lung damage as a cause of admission. Bird lungs are attached to the ventral side of the vertebra and 20%–33% of the tissue between the ribs [28-30]. On the dorsal side of the body, the bird’s chest had more muscle (pectoral) than the waist (abdominal) to enable flight, while part of the pelvis has no muscles attached [31]. Based on these morphological and anatomical features, the lungs may not suffer serious damage due to collisions. Skull congestion was not significantly associated with collision in this study, which was consistent with previous reports [14, 32]. Skull congestion was likely caused by a direct impact on the head or a restriction of blood flow from the trunk to the head, although it can also be caused by postmortem changes [33]. However, in this study, it was difficult to conclude a definitive cause for skull congestion.

Consistent with previous studies [24, 34], collision was not associated with fractures in the studied birds, unlike trauma, which was another reason of admission. Collision generally resulted in more damage to internal organs than open wounds, and the better body condition, the more severe effect on birds. Although histopathological examination was not performed in this study, kidney discoloration and anterior lobe enlargement was possibly due to rupture of renal blood vessels by blunt external force such as collision. We found that among the causes of admission for wild birds that died during the survey period, birds that collided were more likely to show internal organ abnormalities and a higher anterior to posterior kidney lobe ratio, and tended to have a higher fat score than those that died owing to other causes. These results suggested differences in bird injuries concerning the causes of admission and may contribute to the pursuit of a more appropriate rehabilitation protocol. Treatments focusing on potentially damaged organs, such as supportive care for renal damage for birds that were admitted with collision, should be performed promptly. In human medicine, the treatment of renal damage by collision was based on absolute rest [35, 36]. Thus, any handling or treatment that involve unnecessary movement of the birds should be avoided at the time of admission, and absolute cage rest should be adhered for the birds admitted due to collisions to restore renal function as much as possible.

The major limitation of this study was the small sample size; therefore, the causes of admission may not have significantly affected the 50% survival time. Although the number of rehabilitated wild birds could not be anticipated, long-term studies may guarantee larger sample sizes. Several previous studies considered that some species were prone to collision accidents with buildings [14, 24, 37]. Thus, it might have been necessary to clarify the species-specific findings in necropsies.

## ACKNOWLEDGMENTS

We thank all the staff members of the Friends of the Gyotoku Wild Bird Observatory Non-Profit Organization. We are grateful to all members of the Department of Wildlife Medicine of the Nippon Veterinary and Life Science University of Tokyo Prefecture for providing the survey environment.

## REFERENCES

1. Kerr JT, Currie DJ. Effects of human activity on global extinction risk. Conservation Biology. 1995;9(6):1528–38. doi: https://doi.org/10.1046/j.1523-1739.1995.09061528.x.

2. Erickson WP, Johnson GD, Young Jr DP, editors. Anthropogenic Causes with an Emphasis on Collisions’. Bird Conservation Implementation and Integration in the Americas: Proceedings of the Third International Partners in Flight Conference, March 20-24, 2002, Asilomar, California; 2005: US Department of Agriculture, Forest Service, Pacific Southwest Research Station.

3. Gourlay P, Decors A, Moinet M, Lambert O, Lawson B, Beaudeau F, et al. The potential capacity of French wildlife rescue centres for wild bird disease surveillance. European Journal of Wildlife Research. 2014;60(6):865–73. doi: 10.1007/s10344-014-0853-9.

4. Yanagawa H, Shibuya T. Causes of wild bird mortality in eastern Hokkaido II. Research bulletin of Obihiro University. 1996:251–8.

5. Nishi N. The factor of bird collision with windows and the consideration for countermeasure. Japanese Journal of Zoo and Wildlife Medicine. 2010;15(2):95–100. doi: https://doi.org/10.5686/jjzwm.15.95.

6. Mizuta T, Abe Y. Bird strikes on window observed in Amami Oshima Island, Kagoshima Pref., Japan. Bird Research. 2012;8:A25-A33. doi: https://doi.org/10.11211/birdresearch.8.A25.

7. Yoshino T, Uemura J, Watanabe H, Aizawa K, Endo D, Osa Y, et al. Record of wildlife rescue at Rakuno gakuen university wildlife medical centre WAMC (2003-2010). The Journal of Hokkaido Veterinary Medical Association. 2014;58:123–9.

8. Kazama T. Diagnosis and Medical Treatment of Injured/Diseased Wild Birds, Their Captive Maintenance and Release after Recovery. Journal of the Yamashina Institute for Ornithology. 2004;36(1):72–82. doi: https://doi.org/10.3312/jyio.36.72.

9. Suda O, Okubo T, Kanesaka H, Baba K, Ikeya H, Shibata S, et al. Bird’s ecology and prognosis as seen from a chart summary. WRV News Letter. 2010:3–8.

10. Wildlife protection system and hunting law, (1996).

11. Basilio LG, Moreno DJ, Piratelli AJ. Main causes of bird-window collisions: a review. Anais da Academia Brasileira de Ciencias. 2020;92(1). doi: https://doi.org/10.1590/0001-3765202020180745.

12. Hager SB, Trudell H, McKay KJ, Crandall SM, Mayer L. Bird density and mortality at windows. The Wilson Journal of Ornithology. 2008;120(3):550–64. doi: https://doi.org/10.1676/07-075.1.

13. Rodríguez B, Rodríguez A, Siverio F, Siverio M. Causes of raptor admissions to a wildlife rehabilitation center in Tenerife (Canary Islands). Journal of Raptor Research. 2010;44(1):30–9. doi: https://doi.org/10.3356/JRR-09-40.1.

14. Klem D. Bird injuries, cause of death, and recuperation from collisions with windows. Journal of Field Ornithology. 1990;61(1):115–9.

15. Klem D. Avian mortality at windows: the second largest human source of bird mortality on Earth. Tundra to tropics: connecting birds, habitats and people [FULL TEXT]. 2008.

16. Martin GR. Understanding bird collisions with man-made objects: a sensory ecology approach. Ibis. 2011;153(2):239–54.

17. Rebolo-Ifrán N, Di Virgilio A, Lambertucci SA. Drivers of bird-window collisions in southern South America: a two-scale assessment applying citizen science. Scientific reports. 2019;9(1):1–10.

18. Abri B, Vahdati SS, Paknezhad S, Alizadeh S. Blunt abdominal trauma and organ damage and its prognosis. Journal of Research in Clinical Medicine. 2016;4(4):228–32.

19. Beer J. The attempted rehabilitation of oiled sea birds. Wildfowl. 1968;19(19):120–4.

20. Cousins RA, Battley PF, Gartrell BD, Powlesland RG. Impact injuries and probability of survival in a large semiurban endemic pigeon in New Zealand, Hemiphaga novaeseelandiae. Journal of Wildlife Diseases. 2012;48(3):567–74.

21. Ushine N, Sato T, Kato T, Hayama S-i. Analysis of body mass changes in the Black-Headed Gull (Larus ridibundus) during the winter. Journal of Veterinary Medical Science. 2017:17-0099.

22. Sheehy S, Kelly TC, Bourke P, O’Callaghan M, Fennessy GJ, R B. A comparison of the injury syndromes associated with different sources of avian mortality. International Bird Strike Committee; Warsaw2003.

23. Jovani R, Montalvo T, Sabaté S. Fault bars and bacterial infection. Journal of ornithology. 2014;155(3):819–23. doi: http://doi.org/10.1007/s10336-014-1054-8.

24. Hattori K, Kajigaya H. Analysis of wild bird glass collision carcasses. WRV News Letter. 2011:2–7.

25. Lierz M. Avian renal disease: pathogenesis, diagnosis, and therapy. The veterinary clinics of North America Exotic animal practice. 2003;6(1):29-55, v. doi: https://doi.org/10.1016/S1094-9194(02)00029-4.

26. Bartsch R, McConnell E, Imes G, Schmidt J. A review of exertional rhabdomyolysis in wild and domestic animals and man. Veterinary Pathology. 1977;14(4):314–24.

27. Kim SH, Han MC, Kim S, Lee JS. Acute Renal Failure Secondary to Rhabdomyolysis. Acta Radiologica. 1992;33(6):573–6. doi: 10.1080/02841859209173216.

28. Brocklehurst RJ, Schachner ER, Codd JR, Sellers WI. Respiratory evolution in archosaurs. Philosophical Transactions of the Royal Society B. 2020;375(1793):20190140. doi: https://doi.org/10.1098/rstb.2019.0140.

29. Duncker H-R. Structure of avian lungs. Respiration physiology. 1972;14(1-2):44–63. doi: https://doi.org/10.1016/0034-5687(72)90016-3.

30. Maina JN, Nathaniel C. A qualitative and quantitative study of the lung of an ostrich, Struthio camelus. Journal of Experimental Biology. 2001;204(13):2313–30.

31. Suzuki D, Chiba K, VanBuren CS, Ohashi T. The appendicular anatomy of the elegant crested tinamou (Eudromia elegans): Kitakyushu Museum of Natural History and Human History; 2014.

32. de Matos R, Morrisey JK. Emergency and critical care of small psittacines and passerines. Seminars in Avian and Exotic Pet Medicine. 2005;14(2):90–105. doi: https://doi.org/10.1053/j.saep.2005.04.004.

33. Ressel L, Hetzel U, Ricci E. Blunt force trauma in veterinary forensic pathology. Veterinary pathology. 2016;53(5):941–61. doi: https://doi.org/10.1177/0300985816653988.

34. Aymí R, González Y, López T, Gordo O. Bird-window collisions in a city on the Iberian Mediterranean coast during autumn migration. Revista Catalana d’Ornitologia. 2017;33:17–28.

35. Coccolini F, Moore EE, Kluger Y, Biffl W, Leppaniemi A, Matsumura Y, et al. Kidney and uro-trauma: WSES-AAST guidelines. World journal of emergency surgery. 2019;14(1):1–25.

36. Deepak J, Khanday ZS, Bagdi R, Balagopal S, Agarwal P, Madhu R, et al. Three cases of blunt renal trauma in children. Sri ramachandra. 2007:62.

37. Snyder L. Tunnel fliers” and window fatalities. Condor. 1946;48(6):278.

